# Functional T cells trapped behind a stromal wall: a Brake-with-Wall phenotype redefines pancreatic adenocarcinoma immunotherapy resistance

**DOI:** 10.64898/2026.05.11.724216

**Authors:** Jeong-Soon Yong

**Affiliations:** Korea University, Seoul, Republic of Korea

**Keywords:** pancreatic adenocarcinoma, immune evasion, KCNA3, Kv1.3, FAP, cancer-associated fibroblast, T cell exhaustion, immune checkpoint blockade, computational oncology, TCGA

## Abstract

**Background:** Pancreatic ductal adenocarcinoma (PDAC) is the paradigmatic immunotherapy-refractory cancer, with a 5-year survival of approximately 12% and minimal benefit from immune checkpoint blockade (ICB). The dominant mechanistic explanation classifies PDAC as a T cell-excluded “cold” tumor, implying that no functional anti-tumor T cells are available for checkpoint release. Whether this Block-strategy view is correct has not been re-examined under integrated evasion-framework analysis.

**Methods:** We applied a previously developed 16-module immune evasion framework to TCGA-PAAD (n=183), integrated with hub-cytokine analysis (IL-10/TGF-β), Kv1.3-immune channelome data, and clinical trial mapping (12,007 trials). Single-cell validation used two independent PDAC cohorts retrieved through TISCH2: PAAD_CRA001160 (Peng 2019, 35 samples [24 PDAC + 11 adjacent normal], 57,443 cells) and PAAD_GSE154778 (Lin 2020, 16 samples, 14,953 cells), examined for CD8A, TOX, PRF1, KCNA3, and FAP expression by cell type.

**Results:** PDAC scored highest in CAF Wall (z=0.768) and Platelet Cloak (z=0.663) modules; strategy classification yielded Brake — not Block — driven by a positive KCNA3-survival relationship (HR=0.649, 95% CI 0.43–0.97, p=0.037). Single-cell qualitative analysis of TISCH2 violin plots showed that CD8 exhausted T cells (CD8Tex) carried (i) high CD8A, (ii) the highest TOX expression among annotated cell types, (iii) preserved PRF1, and (iv) high KCNA3 expression. FAP was strongly localized to fibroblasts (peak ∼3.0 vs. <0.5 elsewhere). The pattern was reproduced in the second cohort. The optimal three-module attack (MHC restoration + CAF disruption + VEGF blockade) suppressed 10 of 16 evasion modules in silico (62.5%); zero of 370 PDAC immunotherapy trials test this combination.

**Conclusions:** PDAC may not be T cell-cold but T cell-trapped: CD8 T cells with intact Kv1.3 channels appear immobilized behind a FAP-positive cancer-associated fibroblast wall. ICB monotherapy is mechanistically insufficient because the brake is engaged on T cells that cannot reach the tumor. The framework predicts that triple-targeted intervention — checkpoint release + CAF wall disruption + vascular normalization — is the minimum effective strategy. This is a hypothesis-generating computational analysis; prospective experimental and clinical validation are required.

## Background

Pancreatic ductal adenocarcinoma (PDAC) remains one of the most lethal solid malignancies, with a 5-year overall survival of approximately 12% [11]. Unlike melanoma or non-small cell lung cancer, where immune checkpoint blockade (ICB) has produced durable responses, PDAC has shown minimal benefit from anti-PD-1, anti-PD-L1, or anti-CTLA-4 monotherapy. Multiple large trials have reported response rates below 5% in unselected PDAC [13], and the few responders cluster in microsatellite-instability-high subgroups representing 1–2% of all PDAC.

The dominant mechanistic explanation classifies PDAC as a T cell-excluded “cold” tumor. Under this Block-strategy view, the failure of ICB is parsimoniously attributed to the absence of tumor-infiltrating T cells: if no T cells reach the tumor, releasing immune checkpoints has no effect. This has motivated approaches focused on T cell recruitment (chemokines, vaccines) and physical infiltration barriers (CAF disruption, ECM remodeling) [12], rather than on T cell function modulation.

We have recently developed an integrated framework that decomposes tumor immune evasion into 16 mechanistically distinct modules and four operational strategies (Block, Brake, Suppress, Hide), validated across 9,888 TCGA tumors of 33 cancer types [3]; the four-strategy classification was previously evaluated across 31 evaluable cancer types after applying a minimum sample-size threshold [1]. In parallel, we showed that KCNA3 (Kv1.3 voltage-gated potassium channel) is the sole immune-cell-specific ion channel biomarker in pan-cancer bulk RNA sequencing [4], and that IL-10 and TGF-β operate as dual keystone hubs coordinating all 16 evasion modules [2]. Clinical trial mapping further revealed that 60% of 12,007 cancer immunotherapy trials target a single module (checkpoint hijacking), and that combinations targeting more than two modules are exceptionally rare [5].

In this work we apply the integrated framework to PDAC and arrive at a counter-intuitive conclusion: PDAC is not Block but Brake. T cells appear to be present, KCNA3 channels are functional, and the cytotoxic machinery (PRF1) is preserved — yet T cells are immobilized behind a cancer-associated fibroblast wall. We term this phenotype Brake-with-Wall. The clinical consequence is that ICB monotherapy is mechanistically insufficient: it releases a brake on T cells that cannot reach the tumor. The framework predicts a minimum three-module attack — checkpoint release plus CAF wall disruption plus vascular normalization — that is currently tested in zero of 370 registered PDAC immunotherapy trials [5].

## Methods

### Bulk transcriptomic data

TCGA-PAAD RNA-sequencing data (n=183 primary tumors) were obtained from UCSC Xena Pan-Cancer Atlas [10]. We did not perform additional histology-based filtering beyond the primary-tumor restriction provided by Xena, and we acknowledge that prior independent analyses have noted heterogeneity in TCGA-PAAD histology. Sixteen evasion-module signatures and four hub regulators (TGF-β, HIF1α, STAT3, IL-6) were defined and scored as previously described [3], using directionally adjusted gene-level z-scores.

### Strategy classification

Patients were classified into four strategies (Block, Brake, Suppress, Hide) based on T cell infiltration (CD8A, GZMB), checkpoint engagement (PDCD1, CD274, CTLA4), and MHC class I status (HLA-A, HLA-B, B2M), as defined in our prior work [1]. Briefly, T cell-low samples were assigned Block; among T cell-high samples, MHC-low were assigned Hide, checkpoint-high were assigned Brake, and the remainder were assigned Suppress.

### Single-cell validation

Two independent PDAC single-cell cohorts were analyzed via the TISCH2 web portal [6]: PAAD_CRA001160 (Peng et al. 2019 [7], GSA accession CRA001160) and PAAD_GSE154778 (Lin et al. 2020 [8], GEO accession GSE154778). The original Peng study reported 24 PDAC samples and 11 adjacent-normal pancreatic tissues; the original Lin study reported a mixed primary–metastatic PDAC cohort. Cell counts cited in this work (57,443 and 14,953, respectively) reflect the filtered, quality-controlled, and re-annotated cell sets as displayed in the TISCH2 dataset pages on May 9, 2026, and may differ from the original publication counts due to TISCH2’s uniform processing pipeline. Cell type annotations were used as provided by TISCH2.

Five marker genes were examined for expression by cell type using TISCH2-rendered violin plots: CD8A (T cell presence), TOX (exhaustion programming), PRF1 (cytotoxic function), KCNA3 (Kv1.3 channel), and FAP (cancer-associated fibroblast). Cross-cell-type comparisons in this study are qualitative descriptions of TISCH2-rendered violin density, not quantitative tests; raw count re-analysis with formal statistical contrasts is identified as a near-term next step.

### In silico perturbation

Three-module combinations were evaluated by greedy forward selection on TCGA-PAAD samples (n=183). At each step, the module whose knockdown produced the largest mean reduction across the remaining 15 modules was selected. Knockdown was simulated by reducing the log2 expression of all module member genes by a magnitude corresponding to 90% mRNA suppression, with module scores recomputed using baseline z-score parameters [3].

### Clinical trial mapping

PDAC immunotherapy trials were retrieved from ClinicalTrials.gov (API v2) using condition terms (“pancreatic”, “PDAC”, “pancreatic adenocarcinoma”) combined with a curated list of immunotherapy drug names. Each trial was mapped to one or more of the 16 evasion modules via primary mechanism annotation, as detailed previously [5].

### Statistical analysis

Cox proportional hazards regression was performed using the lifelines Python package; hazard ratios and 95% confidence intervals were computed from standardized expression. Survival analysis used Liu et al. integrated TCGA pan-cancer clinical data resource [9]. All analyses used Python 3.12 with pandas, numpy, scipy, scikit-learn, and lifelines.

## Results

### PDAC evasion portfolio is dominated by stromal modules

Among 16 evasion modules scored in TCGA-PAAD (n=183), CAF Wall (H06; mean z=0.768) ranked first and Platelet Cloak (H12; mean z=0.663) ranked second (Fig. 1A). These two stromal modules together contributed more to the total evasion score than the canonical Checkpoint Hijack module (H03; z=0.212), which ranked outside the top five. Treg/M2 Bribery (H02; z=0.498) and Exosome Disruption (H05; z=0.496) ranked third and fourth, respectively.

**Figure 1.**
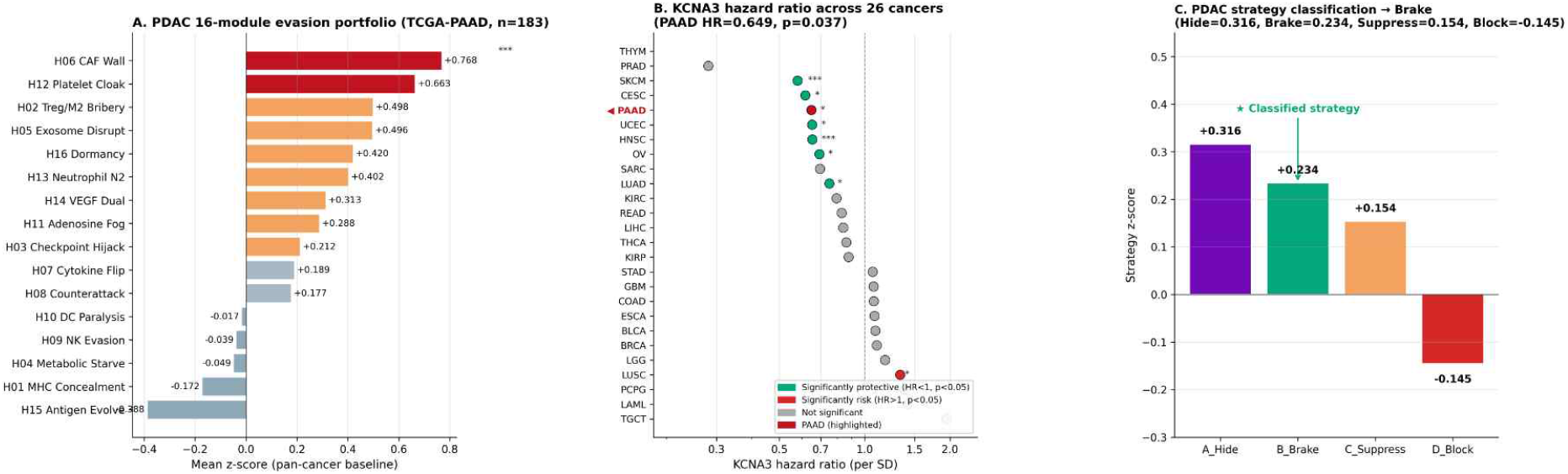
Bulk transcriptomic profile of TCGA-PAAD. (A) Mean z-scores of 16 evasion modules in PDAC (n=183), computed against pan-cancer baseline (33 cancer types, n=9,888); CAF Wall (H06, z=0.768) and Platelet Cloak (H12, z=0.663) rank first and second (highlighted in red). (B) KCNA3 hazard ratio per standard deviation across 26 TCGA cancer types with available survival data; PDAC (red, highlighted) shows significantly protective association (HR=0.649, p=0.037), clustering with other immunotherapy-responsive cancers (SKCM, HNSC, CESC, OV, LUAD). LUSC alone shows significant risk association (HR=1.335, p=0.026). Significance: * p<0.05, *** p<0.001. (C) PDAC strategy z-scores across the four-strategy classification (Hide 0.316, Brake 0.234, Suppress 0.154, Block –0.145). PDAC is assigned the Brake strategy by cascade rule (see Methods 2.2): T cell-high samples that are not MHC-low are evaluated for checkpoint engagement; Brake assignment indicates checkpoint-engaged T cells, not maximum z-score. The classified strategy is highlighted with a star.

This pattern differs from melanoma (SKCM), where Checkpoint Hijack (H03) and MHC Concealment (H01) dominate, and from renal clear cell carcinoma (KIRC), where VEGF (H14) is near-exclusively elevated [3]. The PDAC profile is consistent with a fundamentally stromal, rather than checkpoint-centric, mode of evasion.

### PDAC is classified as Brake, not Block

Strategy classification placed PDAC into the Brake category (Brake-score 0.234, ranked second among four strategy scores: Hide 0.316, Brake 0.234, Suppress 0.154, Block –0.145; assignment by cascade rule defined in Methods, not by maximum score), characterized by T cell presence with checkpoint engagement rather than T cell exclusion (Fig. 1C). This classification was driven primarily by two observations. First, KCNA3 expression — which we previously demonstrated to be immune-cell-specific in pan-cancer single-cell data [4] — was significantly protective in TCGA-PAAD survival analysis (HR=0.649, 95% CI 0.43–0.97, p=0.037; Fig. 1B), consistent with the presence of functionally competent immune cells. Second, TGFB1 was not significantly associated with survival in PDAC (HR=1.055, p=0.80), in contrast to colon and renal cancers where TGFB1 marks fibroblast-driven exclusion [2,15,16].

### Single-cell qualitative observation suggests functional T cells in PDAC

Examination of PAAD_CRA001160 (TISCH2-processed; 57,443 cells; Fig. 2) revealed an annotated CD8Tex (CD8 exhausted T cell) cluster, with B cells, plasma cells, monocytes/macrophages, dendritic cells, fibroblasts, malignant ductal cells, and stromal compartments also represented. By visual inspection of violin plots, CD8A expression was prominent in CD8Tex (and B cell) clusters, qualitatively consistent with the presence of CD8+ T cells in primary PDAC tumors (Fig. 2A). TOX, the exhaustion master transcription factor, showed the highest violin density among annotated cell types in this cohort (Fig. 2B). Critically, PRF1 (perforin) — the principal cytotoxic effector — was visibly preserved in CD8Tex relative to non-lymphoid clusters (Fig. 2C), suggesting that the killing machinery is at least partially intact despite the exhaustion phenotype.

**Figure 2.**
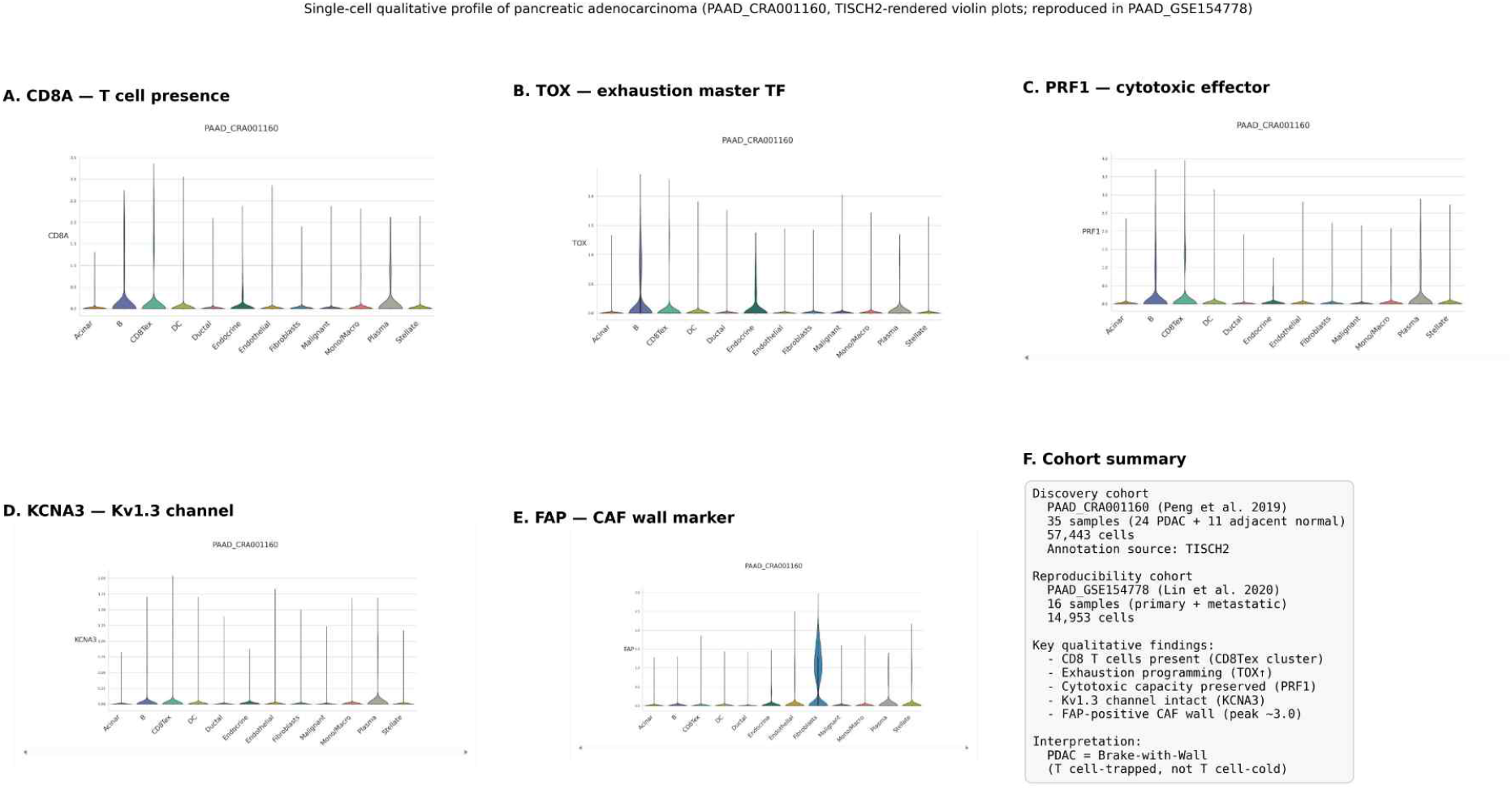
Single-cell qualitative profile of pancreatic adenocarcinoma (PAAD_CRA001160; Peng 2019; n=35 samples, 24 PDAC + 11 adjacent normal; 57,443 cells; TISCH2-processed). Violin plots of cell-type-specific expression for (A) CD8A, marker of CD8 T cell presence; (B) TOX, master regulator of T cell exhaustion; (C) PRF1, principal cytotoxic effector; (D) KCNA3, encoding the Kv1.3 voltage-gated potassium channel; (E) FAP, marker of cancer-associated fibroblasts. (F) Cohort summary. Violin plots are reproduced from the TISCH2 web portal.

KCNA3 expression was concentrated in the CD8Tex cluster, with smaller contributions from B and Mono/Macro cells, and near-absence in malignant, ductal, fibroblast, and endothelial compartments (Fig. 2D). This single-cell pattern is qualitatively consistent with the previously reported pan-cancer immune-specific localization of KCNA3 [4,14] and supports a functional Kv1.3 channel in the resident PDAC T cell pool.

### FAP-positive fibroblasts form a dense stromal wall

In the same PAAD_CRA001160 dataset, FAP expression was strongly localized to the Fibroblast cluster, with violin density peaking near 3.0 (log-normalized) and median expression substantially above zero — the highest of any annotated cell type (Fig. 2E).

All non-fibroblast clusters (including malignant cells, T cells, and endothelial cells) showed peak FAP expression below 0.5 with median near zero. This sharp localization is qualitatively consistent with the presence of a FAP-positive cancer-associated fibroblast wall [12,17], the molecular substrate hypothesized to underlie the H06 CAF Wall module score.

### Independent cohort reproduces the pattern

To assess reproducibility, we examined PAAD_GSE154778 (TISCH2-processed; 14,953 cells; mixed primary and metastatic) using the same five marker genes. KCNA3 expression was again concentrated in CD8 T cells (highest among lymphoid clusters) and Mono/Macro cells, with negligible expression in epithelial, malignant, and fibroblast compartments. FAP was again strongly localized to fibroblasts (peak ∼2.87 vs. <0.5 elsewhere). The two independent cohorts thus yielded qualitatively concordant patterns at the level of cell-type-specific expression, supporting the generalizability of the Brake-with-Wall pattern across PDAC.

### The minimum effective combination is untested

Greedy forward selection of three-module knockdowns identified MHC restoration (H01) + CAF Wall disruption (H06) + VEGF blockade (H14) as the minimum effective combination for PDAC, suppressing 10 of 16 evasion modules in silico (62.5%). Of 370 registered PDAC immunotherapy clinical trials retrieved from ClinicalTrials.gov [5], none test this three-module combination. The closest tested approach — chemotherapy (gemcitabine, FOLFIRINOX) plus anti-PD-L1 — does not include either MHC restoration (e.g., hypomethylating agents) or CAF wall disruption (e.g., FAP-directed agents). Trials of FAP-directed approaches (FAP-2286, sibrotuzumab, FAP-targeted CAR-T) remain in early phase, with fewer than ten registered PDAC trials each [5].

## Discussion

### Reframing PDAC: T cell-trapped, not T cell-cold

The dominant clinical narrative classifies PDAC as immunologically cold and concludes that ICB has no biological substrate to act on. Our analysis suggests this framing is incomplete. PDAC tumors do contain CD8 T cells with the molecular features of recently activated, exhausted-but-not-terminal effectors: high TOX (exhaustion) coexists with preserved PRF1 (cytotoxic capacity) and KCNA3 (Kv1.3 channel function). The mechanistic problem is therefore not absence of immune cells but spatial confinement: a FAP-positive CAF wall and a platelet/VEGF-driven vascular barrier together immobilize an otherwise competent T cell pool. We propose Brake-with-Wall as a more precise descriptor than Block.

### Implications for immune checkpoint blockade

If PDAC T cells are present but trapped, ICB monotherapy releases a brake on cells that cannot reach the tumor. This may explain the consistently low ICB response rates in PDAC despite measurable PD-L1 expression in some patients [13]. The framework predicts that ICB efficacy would improve substantially under combination with CAF wall disruption (FAP-directed agents, anti-FAP-TGFβ bispecifics, LRRC15-directed antibody-drug conjugates [17]) and vascular normalization (bevacizumab, ramucirumab), with optional MHC restoration (decitabine, azacitidine) for the subset of tumors with measurable HLA loss.

### KCNA3 as a candidate biomarker

The protective KCNA3 hazard ratio in TCGA-PAAD (HR=0.649, p=0.037) is, on its face, modest in effect size and not corrected for multiple testing across 33 cancer types [4]. However, when interpreted alongside the qualitative single-cell finding that KCNA3 is concentrated in CD8Tex cells, the bulk signal acquires biological meaning: KCNA3-high PDAC tumors may harbor more functionally competent T cells. This is consistent with the pan-cancer pattern reported previously [4] and motivates KCNA3 as a candidate biomarker for selecting PDAC patients into trials of T cell-augmenting agents (anti-PD-1, Kv1.3 openers, IL-2 variants) combined with stromal-directed therapy.

### Why the framework-recommended combination has not been tested

The MHC + CAF + VEGF triple combination is absent from the 370 PDAC immunotherapy trials in our dataset for three reasons. First, FAP-directed agents remain in early development with limited clinical access. Second, three-drug combinations face compounding regulatory, intellectual property, and safety hurdles not faced by two-drug combinations. Third, the prevailing Block-strategy view of PDAC has directed trial investment toward T cell recruitment (vaccines, oncolytic viruses, CAR-T) rather than T cell unblocking. The framework reframes this priority [5].

### Limitations

This is a computational, hypothesis-generating analysis. Several limitations warrant explicit acknowledgement. First, the strategy classification, module scoring, and in silico perturbation are derived from bulk TCGA RNA-seq with previously published gene panels [1,3]; they have not been validated experimentally in PDAC tissue or PDAC-specific patient-derived models. Second, the single-cell observations are qualitative descriptions of TISCH2-rendered violin plots [6]; we did not perform raw count re-analysis with formal statistical contrasts. Quantitative claims such as “highest expression among cell types” should be read as visual-density observations and require quantitative confirmation. Third, cell counts cited (57,443 and 14,953) reflect TISCH2’s filtered/annotated outputs and may differ from the source publications’ counts. Fourth, the four-strategy framework was defined in our prior work [1] and remains to be externally validated by independent groups. Fifth, the clinical trial gap analysis [5] depends on drug-to-module mapping accuracy estimated at approximately 80%; the underlying registry is also U.S./EU-weighted. Sixth, hazard ratios derived from in silico perturbation are not estimates of expected clinical effect sizes. Seventh, TCGA-PAAD is known to contain a fraction of histologies that may not be PDAC; we did not apply additional histology-based filtering. Prospective validation in clinical cohorts and wet-lab model systems is essential.

## Conclusions

Pancreatic ductal adenocarcinoma may not be a T cell-excluded tumor but a T cell-trapped one. CD8 T cells with qualitatively preserved cytotoxic machinery (PRF1) and intact Kv1.3 channel (KCNA3) appear to reside within PDAC tumors but are immobilized behind a FAP-positive cancer-associated fibroblast wall. The framework predicts that the minimum effective intervention is a three-module combination — MHC restoration plus CAF wall disruption plus VEGF blockade — that is currently absent from the global PDAC clinical trial portfolio. KCNA3 is a candidate biomarker for selecting PDAC patients into combination trials. The reframing from Block to Brake-with-Wall has direct implications for trial design and combination prioritization in PDAC immunotherapy, pending prospective experimental and clinical validation.

## Abbreviations

CAF: cancer-associated fibroblast
CD8Tex: CD8 exhausted T cell
FAP: fibroblast activation protein
HR: hazard ratio
ICB: immune checkpoint blockade
KCNA3: potassium voltage-gated channel subfamily A member 3 (Kv1.3)
MHC: major histocompatibility complex
PDAC: pancreatic ductal adenocarcinoma
PD-1: programmed cell death protein 1
PD-L1: programmed death-ligand 1
PRF1: perforin 1
TCGA: The Cancer Genome Atlas
TGF-β: transforming growth factor beta
TISCH2: Tumor Immune Single-cell Hub 2
TOX: thymocyte selection associated HMG box
VEGF: vascular endothelial growth factor

## Declarations

### Ethics approval and consent to participate

This study used only publicly available, de-identified datasets (TCGA via UCSC Xena, ClinicalTrials.gov API v2, TISCH2 web portal). No human subjects were recruited, no human tissue was collected, and no animals were used. Institutional review board approval was not required as all data were previously published and publicly accessible in accordance with their original ethics approvals.

### Consent for publication

Not applicable.

### Availability of data and materials

All data analyzed in this study are publicly available. TCGA-PAAD bulk RNA-seq and clinical data are available from UCSC Xena (https://xenabrowser.net). Clinical trial data are available from ClinicalTrials.gov (https://clinicaltrials.gov/api/v2). Single-cell RNA-seq data are available through the TISCH2 web portal (http://tisch.comp-genomics.org), which curates and uniformly reprocesses datasets originally deposited at: GSA accession CRA001160 (Peng et al. 2019) and GEO accession GSE154778 (Lin et al. 2020). Analysis code is available from the corresponding author upon reasonable request.

### Authors’ contributions

J.-S.Y. conceived the study, performed all analyses, and wrote the manuscript. The author has read and approved the final manuscript.

## Acknowledgements

Not applicable.

## Use of artificial intelligence

AI assistance (Claude, Anthropic) was used for code generation, data analysis, and manuscript drafting. The AI did not meet authorship criteria and is not listed as an author. The corresponding author assumes full responsibility for all scientific content, analytical decisions, and conclusions.

